# Inhalation frequency controls reformatting of mitral/tufted cell odor representations in the olfactory bulb

**DOI:** 10.1101/242784

**Authors:** Marta Díaz-Quesada, Isaac A. Youngstrom, Yusuke Tsuno, Kyle R. Hansen, Michael N. Economo, Matt Wachowiak

## Abstract

In mammals olfactory sensation depends on inhalation, which controls activation of sensory neurons and temporal patterning of central activity. Odor representations by mitral and tufted (MT) cells, the main output from the olfactory bulb (OB), reflect sensory input as well as excitation and inhibition from OB circuits, which may change as sniff frequency increases. To test the impact of sampling frequency on MT cell odor responses, we obtained whole-cell recordings from MT cells in anesthetized male and female mice while varying inhalation frequency via tracheotomy, allowing comparison of inhalation-linked responses across cells. We characterized frequency effects on MT cell responses during inhalation of air and odorants using inhalation pulses and also ‘playback’ of sniffing recorded from awake mice. Inhalation-linked changes in membrane potential were well-predicted across frequency from linear convolution of 1 Hz responses and, as frequency increased, near-identical temporal responses could emerge from depolarizing, hyperpolarizing or multiphasic MT responses. However, net excitation was not well predicted from 1 Hz responses and varied substantially across MT cells, with some cells increasing and others decreasing in spike rate. As a result, sustained odorant sampling at higher frequencies led to increasing decorrelation of the MT cell population response pattern over time. Bulk activation of sensory inputs by optogenetic stimulation affected MT cells more uniformly across frequency, suggesting that frequency-dependent decorrelation emerges from odor-specific patterns of activity in the OB network. These results suggest that sampling behavior alone can reformat early sensory representations, possibly to optimize sensory perception during repeated sampling.

**Significance statement:** Olfactory sensation in mammals depends on inhalation, which increases in frequency during active sampling of olfactory stimuli. We asked how inhalation frequency can shape the neural coding of odor information by recording from projection neurons of the olfactory bulb while artificially varying odor sampling frequency in the anesthetized mouse. We found that sampling an odor at higher frequencies led to diverse changes in net responsiveness, as measured by action potential output, that were not predicted from low-frequency responses. These changes led to a reorganization of the pattern of neural activity evoked by a given odorant that occurred preferentially during sustained, high-frequency inhalation. These results point to a novel mechanism for modulating early sensory representations solely as a function of sampling behavior.

## Introduction

A fundamental step in understanding sensation is determining how neural circuits in the brain transform sensory inputs into meaningful patterns of activity among central neurons. In all sensory modalities, the detection and initial encoding of sensory information is a dynamic process that can be actively regulated by sampling behavior. Understanding how central circuits process dynamic sensory inputs in the context of active sampling is thus critical for understanding the neural basis of sensation.

In olfaction, a primary sensory modality in many mammals, olfactory sensation depends on the inhalation of air through the nasal cavity. Inhalation determines the initial temporal structure of sensory input to the brain and is a major determinant of neural dynamics at all subsequent processing stages (Macrides and Chorover, 1972; Onoda and Mori, 1980; Sobel and Tank, 1993; Kepecs et al., 2006; Schaefer and Margrie, 2007; Wachowiak, 2011). Odorant inhalation is actively and precisely controlled during wakefulness, with animals showing distinct repertoires of active sampling (‘sniffing’) that likely reflect strategies for optimally extracting and processing sensory information depending on task demand (Laing, 1983; Kepecs et al., 2007; Wesson et al., 2008; Wachowiak, 2011). One prominent strategy involves sustained increases in inhalation frequency; such high-frequency sniff bouts are a hallmark of exploratory behavior in rodents and other mammals (Welker, 1964; Macrides, 1975). High-frequency sniffing elicits partial adaptation of olfactory sensory neuron inputs to the olfactory bulb (OB), which may increase the salience of newly-encountered odorants relative to background stimuli (Verhagen et al., 2007). However, how active sampling impacts the central processing of olfactory information is less clear.

Circuits within the OB have long been hypothesized to differentially process OSN inputs occurring across the behavioral range of sniff frequencies (Wachowiak and Shipley, 2006). For example, OB slice experiments have shown that increasing input frequency can alter the relative strength and temporal dynamics of excitation and inhibition within the OB (Young and Wilson, 1999; Balu et al., 2004; Schoppa, 2006). Two general predictions from these studies are that high-frequency sniffing increases the reliability and temporal precision of MT cell firing relative to the respiratory cycle and enhances the signal-to-noise ratio of odorant-evoked MT cell responses (Balu et al., 2004; Hayar et al., 2004; Wachowiak and Shipley, 2006; Shao et al., 2013). In vivo tests of these hypotheses have so far consisted of extracellular MT cell recordings and have reported conflicting results, with some studies reporting a temporal sharpening of MT cell responses and reduced MT cell excitation (Bathellier et al., 2008; Carey and Wachowiak, 2011) and others reporting response patterns that are consistent with a simple linear extrapolation of unitary, low-frequency responses (Gupta et al., 2015).

Here, we investigated how inhalation frequency shapes MT cell membrane potential and spiking responses in vivo using whole-cell current clamp recordings in anesthetized mice. We varied inhalation frequency using a paradigm that allowed precise comparison of inhalation-linked response patterns across frequency and across different recordings. We found that inhalation-linked temporal patterns of membrane potential changes were generally well predicted by linear summation of low-frequency responses in absolute time rather than inhalation phase. However, net excitation as measured by MT spike output was not well predicted from low-frequency responses; instead, increasing inhalation frequency had diverse effects on excitability among different MT cells that could not be ascribed to intrinsic differences across cells or cell types. We show that these results predict that odor representations across a MT cell population are reformatted during sustained high-frequency sampling of odorant, and that such reformatting does not occur when OSN inputs are activated in bulk by optogenetic stimulation. Overall these results point to a novel mechanism for modulating early odor representations solely as a function of sampling behavior.

## Materials and Methods

### Animals

Experiments were performed on male and female mice ranging in age from 2 - 4 months. Mice used were either wild-type C57/Bl6 or mice from either of two strains: PCdh21-Cre (MMRRC Stock #030952-UCD) (Wachowiak et al., 2013) or OMP-SpH (Jax Stock #004946) (Bozza et al., 2004), both in the C57/Bl6 background. For optical stimulation experiments, the OMP-ChR2-YFP line (Smear et al., 2011) was used. OMP-ChR2-YFP mice were a gift of T. Bozza, Northwestern University. Both of the OMP strains were used as heterozygous for the OMP knockin in all experiments. All procedures were carried out following National Institutes of Health Guide for the Care and Use of Laboratory Animals and were approved by the University of Utah Institutional Animal Care and Use Committee.

### Whole-cell recordings

Mice were anesthetized with pentobarbital (50–90 mg/kg) and supplemented as needed throughout the experiment; in two mice used for the sniff playback experiments (Fig. 6), anesthesia was maintained with isoflurane (0.5–1.25%). Body temperature and heart rate were monitored and maintained at ~37 ºC and ~ 400 beats per minute. A double tracheotomy was performed for artificial inhalation with the mouse breathing freely through one tracheotomy tube and the second tube connected to a solenoid-gated vacuum source (Fig. 1A) or sniff playback device as described previously(Cheung et al., 2009; Carey and Wachowiak, 2011). The dorsal surface of the cranium was exposed and the animal secured with a custom bar cemented to the skull, then a chamber was created with dental cement around the dorsal OB. When OMP-spH mice were used, the overlying bone was thinned and wide-field epifluorescence signals were acquired to confirm odorant-evoked activity in the OB. A small craniotomy (1 × 1 mm) and durectomy was then performed and the exposed OB surface kept immersed in ACSF.

**Figure 1.**
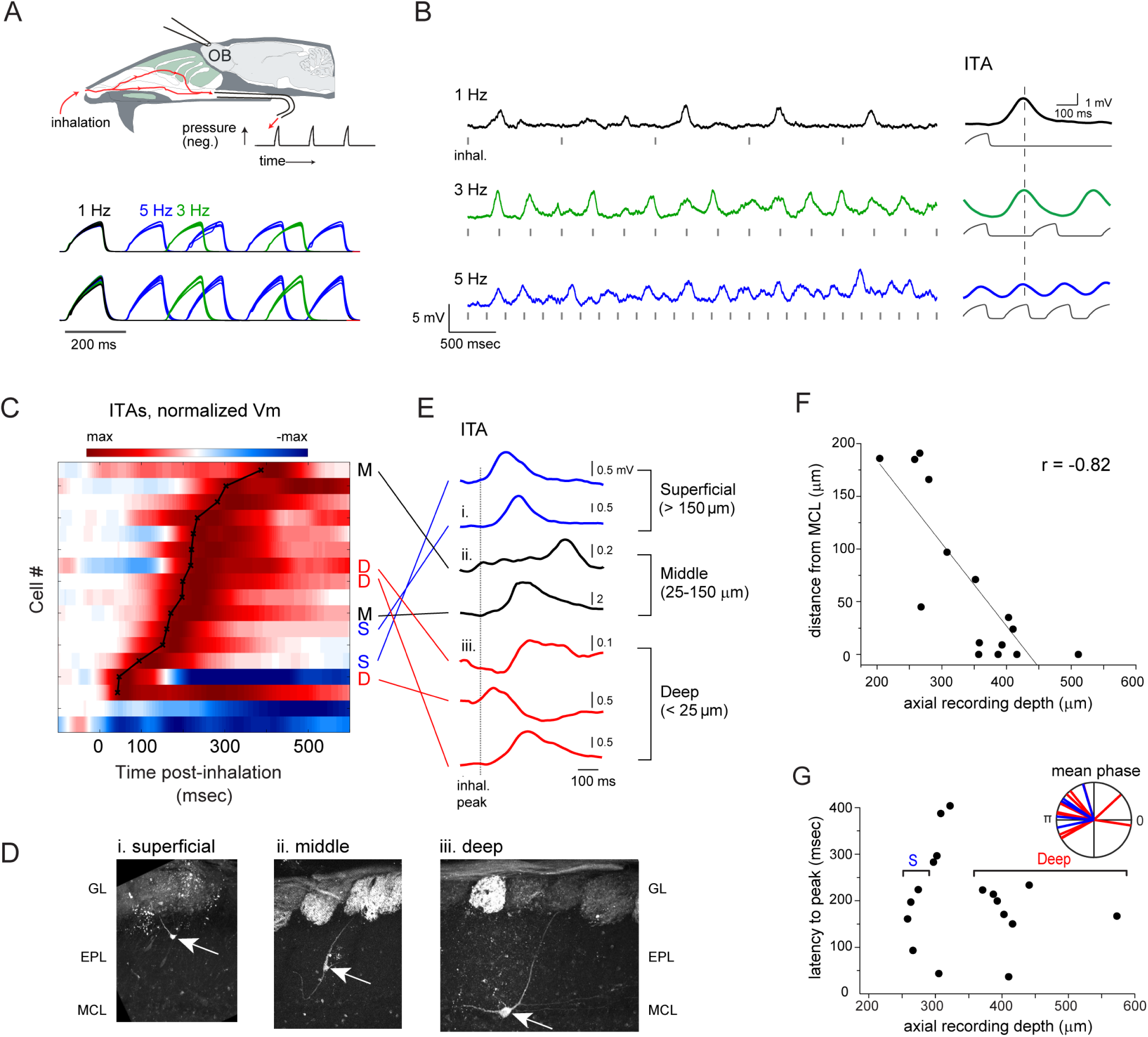
Inhalation-linked modulation of membrane potential in a fraction of MT cells. A. Schematic of artificial inhalation paradigm. Negative pressure transients were applied to the nasopharynx while performing whole-cell recordings from OB neurons. Traces show pressure transients measured in parallel with each recording, delivered at 1, 3 and 5 Hz, from two different recordings. Negative pressure (inhalation) is upward. Each set of traces includes overlays of 18–29 trials. B. Membrane potential (V_m_) traces from a superficial MT cell (shown in panel D) during inhalation at 1, 3 and 5 Hz. Hash marks indicate onset of each inhalation, for visual reference. The ‘inhalation-triggered average’ (ITA) V_m_ response to repeated inhalations at each frequency is shown to the right of each trace, along with the simultaneously-measured pressure transient (gray). Vertical dashed line indicates time of peak ITA depolarization at 1 Hz for comparison across frequencies. C. Pseudocolor representation of ITAs for all recorded cells showing significant V_m_ modulation by 1 Hz inhalation (17 of 36 total cells). Cells are ordered in rows and sorted in order of latency to peak depolarization, indicated by overlaid black plot. Pseudocolor range is set to twice the maximum absolute deviation from baseline V_m_ (max to -max), normalized separately for each ITA and displayed in 0.1 msec time-bins. Time 0 indicates the time of peak negative pressure, which is 150 msec after inhalation onset. Morphologically identified cells (defined as in panels D – F) are labelled as deep (D), middle (M) or superficial (S) to the right of their corresponding row. D. Examples of superficial, middle and deep MT cells after partial morphological recovery. Arrows indicate soma, with primary dendrite extending into a glomerulus. GL, glomerular layer; EPL, external plexiform layer; MCL, mitral cell layer. Fluorescence in glomeruli is from synaptopHluorin expressed in OSNs in OMP-spH mice (see Text). E. ITA Vm traces for each of the 7 identified cells in C, classified by depth from superficial to deep. Traces begin at inhalation onset; dashed line indicates time of peak inhalation (corresponding to t = 0 in C). F. Plot of soma position relative to MCL versus axial depth of the recording pipette for all filled cells. Line indicates linear fit to data. G. Latency to peak depolarization for all inhalation-modulated cells, measured from the ITA at 1 Hz and plotted as a function of recording depth. Brackets indicate depth range used to classify MT cells as deep or superficial (deep, > 350 µm; superficial, <275 µm). *Inset*, vectors representing the mean phase of the Vm ITA with respect to the inhalation cycle from superficial (blue, n = 5) and deep (red, n = 11) MT cells significantly modulated by 1 Hz respiration (p<0.05, Rayleigh’s test).

Whole-cell recordings were made with glass electrodes (4.3–7 MΩ) drawn on a horizontal puller (P97; Sutter Instruments) and filled with the following internal solution (in mM): 120 K-gluconate, 20 KCl, 10 HEPES, 7 diTrisPhCr, 4 Na2ATP, 2 MgCl2, 0.3 Tris-GTP, 0.2 EGTA, or 135 K-gluconate, 4 KCl, 10 HEPES, 10 phosphocreatine, 4 MgATP, 0.3 GTP, both buffered to pH 7.3 with KOH (Sigma-Aldrich) and 285–310 mOsm. In some experiments, Alexa594 (100 μM) or sulforhodamine 101 (20 μM) (Invitrogen) were added to the internal solution for morphological recovery. The pipette was advanced axially by 200 - 500 µm into the dorsal OB at an angle of roughly 30 degrees using a piezoelectric micromanipulator (Luigs and Neumann SM-6, Germany). For the sniff playback recordings the pipette was advanced vertically. Positive pressure was maintained on the pipette until electrode resistance increased, then slight negative pressure was used to obtain a gigaohm seal and then suction and/or an electronic buzz were applied to obtain the whole-cell configuration. Recordings were performed in current-clamp mode with a Multiclamp 700A amplifier (Molecular Devices), digitized at 10 kHz, and stored to disk. Input resistance across cells included for analysis (n = 56) had a median input resistance of 125 MΩ (107 MΩ interquartile range). Most recordings lasted between 15 – 60 minutes. To test for baseline drift across trials, we calculated the difference in spike threshold V_m_ and the baseline V_m_ between groups of trials at each inhalation frequency. Resting V_m_ relative to spike threshold V_m_ was -10.5 ± 3.7 mV (n = 912 trials, 36 cells with sufficient spikes). Stimulation and recording protocols were controlled using pClamp10 (Molecular Devices).

For post-hoc morphological recovery of recorded cells, the mouse was deeply anesthetized with an overdose of sodium pentobarbital and transcardially perfused with phosphate-buffered saline (PBS) followed by 4% paraformaldehyde. Heads were left overnight at 4°C before the brain was extracted. Filled cells were visualized directly by Alexa 594 or sulforhodamine fluorescence from 100–200 μm vibratome sections.

### Artificial inhalation and stimulation protocols

Artificial inhalation was controlled by a three-way solenoid that directed vacuum to the nasopharynx via the tracheotomy tube, as described previously (Wachowiak and Cohen, 2001; Spors et al., 2006). The duration of each inhalation pulse was set to 150 ms, resulting in negative pressure transients applied to the nasopharynx with mean ± s.d. half-width of 77 ± 4 msec (Fig. 1A). Inhalation timing and consistency was monitored with a pressure sensor. For experiments, inhalation pulses were repeated at 1, 3 or 5 Hz as described in Results. Baseline inhalation frequency was maintained at 2 Hz.

Odorants were presented as dilutions from saturated vapor (% s.v.) in cleaned, humidified air using a custom olfactometer under computer control (Bozza et al., 2004; Verhagen et al., 2007). Final odorant concentrations ranged from 0.1% to 2% s.v; in some cases, odorants were diluted 1:10 in mineral oil before the vapor phase dilution with the assumption of an identical dilution of the final vapor concentration. Odorants were screened beginning at the lowest concentration possible and this was increased until a clear response (EPSPS, IPSPs or spikes) was detected; we routinely screened several odorants before obtaining a reliable response in a given cell. Each trial was 25 seconds long, with the first 5 sec consisting of inhalation of ambient air at the tested frequency of 1, 3 or 5 Hz, then odor presentation for 10 sec, followed by a 10 sec post-odor period. A given odorant was presented 3 – 6 times at each of the three frequencies, with the different frequencies interleaved in randomized order.

For optical stimulation of olfactory sensory neurons in OMP-ChR2 mice, blue light was directed onto the dorsal OB surface using a 1 mm optical fiber connected to a 470 nm LED and controller (LEDD1B, Thorlabs). Total power emission from the fiber ranged from 65 μW to 6.2 mW. Light pulses were 75 - 100 ms in duration to approximate the duration of inhalation and were applied at 1, 3 and 5 Hz in 15-sec long trains. Each frequency was applied 3–5 times with frequencies interleaved.

### Measurement and playback of sniffing in awake mice

Intranasal pressure was measured from awake, head-fixed mice engaged in a go/no-go odor discrimination task, using procedures described previously (Wesson et al., 2008) (Wachowiak et al., 2013). Pressure waveforms were analyzed from multiple behavioral trials without respect to the discrimination task or odorant identity (numbers of mice and trials analyzed are given in Results). Sniff playback was performed using a custom solenoid-driven device connected to a syringe, described previously, which reproduced recorded pressure transients with high fidelity (Cheung et al., 2009; Carey and Wachowiak, 2011). Three playback traces were constructed: the first was a single inhalation taken during passive, low-frequency (1 - 2 Hz) breathing; this inhalation was repeated at 1 Hz to serve as a baseline for comparison with higher-frequency sniff bouts. The second trace consisted of natural bouts of approximately 5 Hz sniffing repeated several times, while the third consisted of several epochs of ‘exploratory’, often higher-frequency sniffing, (3 - 8 Hz), that better represented the variability in frequency and pressure waveforms of recordings from awake mice (traces are shown in Figure 6B). These traces were used to drive the sniff playback device through custom software written in Labview. The sniff playback solenoid drove a syringe plunger which was connected via teflon tubing to the tracheotomy tube. The syringe size and gain of the playback was chosen to generate air fluxes matching the average tidal volume of an adult mouse (0.2 ml). A pressure sensor connected to the sniff playback line monitored device output, and this signal was used to verify reproducibility and to align all responses during analysis.

### Data analysis

To analyze inhalation-linked patterning of membrane potential (Vm), inhalation-triggered averages (ITAs) of Vm were calculated by first clipping action potentials at 10 mV above the mean Vm for the recording and averaging over a 700 ms time-window around the time of inhalation onset after low-pass filtering at 200 Hz to remove any remaining spike signal from the third inhalation onward. Significant modulation of Vm by inhalation was tested using bootstrapped values as follows: For each cell, the first 5 secs of all trials at 1 Hz inhalation were selected and five traces (corresponding to 5 inhalations in the pre-odor period) the length of the calculated ITAs (700 ms) were randomly chosen from each trial for that cell; for example, one cell with 3 trials would have 15 randomly selected traces. All of these randomly selected traces were averaged together to calculate a randomized bootstrapped ITA. An excitatory and inhibitory amplitude was calculated for each randomized bootstrapped ITA as the difference between the maximum/minimum point for the last 600 ms of the ITA compared to the average of the first 100 ms of the trace. This process was repeated 10,000 times for each cell, calculated excitatory and inhibitory amplitudes were sorted from the bootstrapped calculations and the 95th percentile amplitude was selected as the significance threshold for excitatory and inhibitory responses. This 95th percentile threshold was compared to the excitatory and inhibitory response of the ITA calculated from the recorded trace, during both pre-odor and odorant-evoked ITAs to determine which cells were significantly modulated by inhalation.

For measurement of spike times and spike rates, action potentials were detected from an increase in the first derivative of the Vm and spike times set to the time at which the Vm surpassed the threshold then placed into 10 ms bins before averaging across inhalations. Inhalation triggered spike histograms were calculated from the average number of spikes in each 10 ms bin of a 700 ms window around inhalation onset, averaged across inhalations at a given frequency. Spike raster marks (e.g., Figs. 2E) indicate 10 ms bins with at least one spike during any inhalation. Mean spike rate was calculated as the average number of spikes per 1-sec period and peak spike rate was calculated as the maximum number of spikes within any 200 ms window of each 1-sec period, averaged across trials including at least one spike per 1-sec period. In the optical stimulation experiments, spiking was measured as the mean number of spikes per 200 ms window following each light pulse (i.e., spikes per pulse).

**Figure 2.**
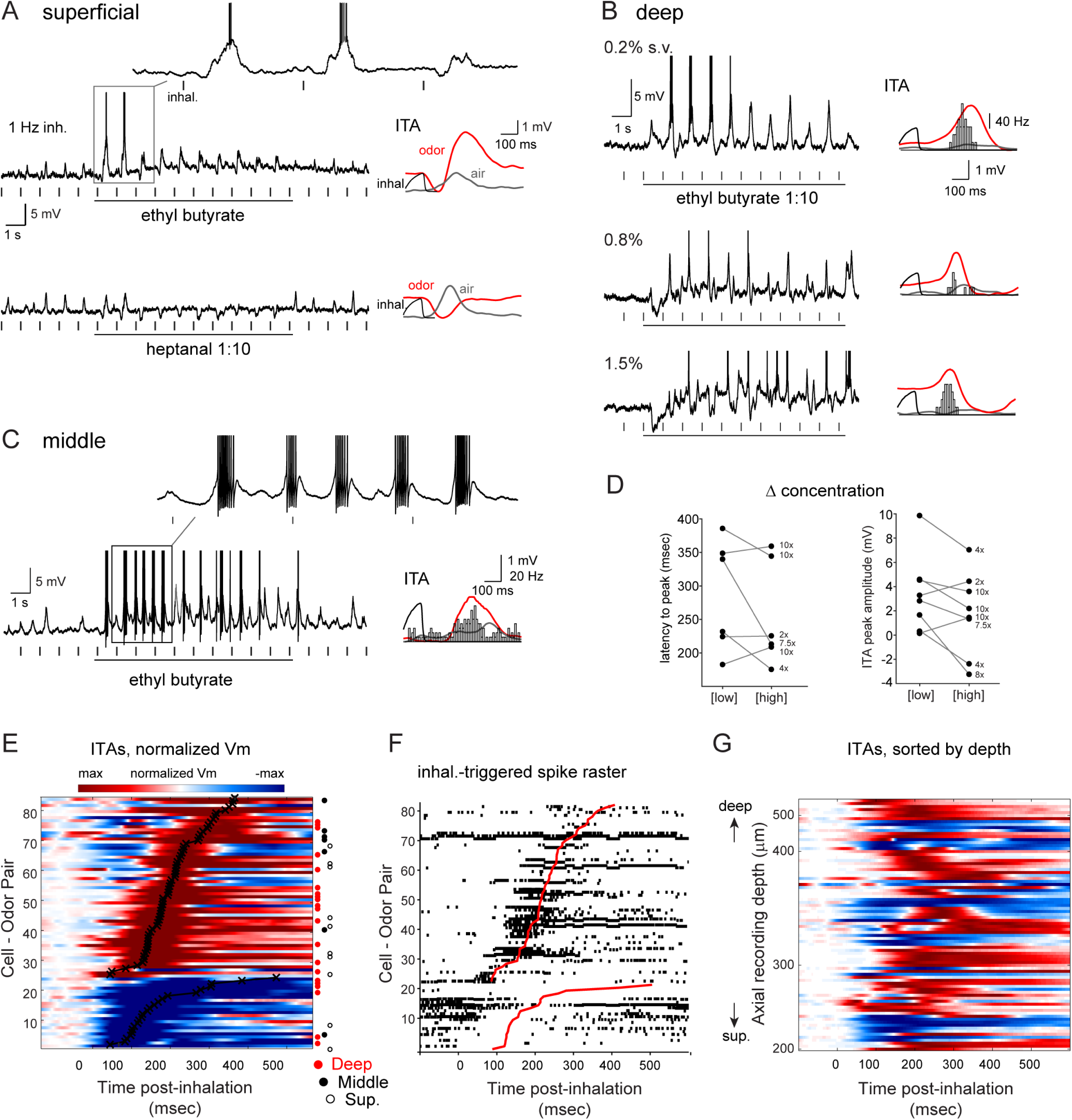
Diverse inhalation-linked temporal patterning of odorant-evoked responses. A - C. Examples from three filled MT cells showing diverse responses to odorant at 1 Hz inhalation. Raw V_m_ traces show response to a single, 10-sec odorant presentation. Plots at right show ITA during odorant inhalation (red) compared to ITA of ambient air prior to odorant presentation (gray). ITAs include spike histograms (bins, 10 msec), generated from seconds 3 - 10 of stimulation. A: Distinct response patterns to two different odorants in a superficial MT cell (same cell as in Fig. 1B). B: Progression of inhalation-driven V_m_ and spiking response patterns with increasing odorant concentration in a deep MT cell. C: Entrainment of odorant-evoked spike bursts to inhalation in a middle MT cell. D. Plot of ITA latency, measured as time-to-peak (left) and amplitude (right) as a function of odorant concentration. Connected points show values for the same MT cell tested at a low or high concentration; numbers to the right of each high concentration point indicate relative increase in concentration. Latency values were measured for depolarizing ITAs only (6 cell-odor pairs); amplitude values were measured for either polarity and so included two additional cell-odor pairs (from 6 MT cells total). E. Pseudocolored time-course of odorant-evoked ITA for all significantly-modulated cell-odor pairs (82 of 90), sorted by latency to peak depolarization or hyperpolarization (lower rows) as indicated by black plot. Approximately 30% of cells show a predominately hyperpolarizing response. Rows corresponding to filled superficial (S, open circle), middle (M, filled circle) or deep (D, red circle) cell-odor pairs are indicated to the right of each row. Time 0 indicates time of peak inhalation. F. Inhalation-linked spike rasters for the same cell-odor pairs as in (E), sorted as in (E). Rasters are compiled across trials. G. Same Vm ITAs as in (E), sorted by axial recording depth instead of latency. Note no clear progression of ITA temporal patterns from superficial to deep recordings.

To test for frequency-dependent effects on inhalation-linked patterning, ITAs of the Vm at 1 Hz were used as a kernel. The first point of the kernel was set to zero before convolving with a vector of sniff onset times at 1,3 and 5 Hz, or in the case of sniff playback trials, the point 100 msec before the peak of each sniff (note that this could lead to kernels with non-zero mean), and then subtracting the experimental mean pre-odor baseline Vm at each frequency from the convolved traces. Experimental and convolved ITAs were compared by measuring the cross correlation following mean subtraction. The maximum cross correlation value (rpeak) and the lag value of that peak were compared across frequencies.

All analyses and statistical tests were performed using custom scripts written in Matlab 2015b (Mathworks) or in the companion package of R version 3.3.2. Comparison of mean phase differences were performed using the Circular Statistics Toolbox (Berens, 2009). All summary statistics are reported as mean ± standard deviation (s.d.) unless noted otherwise. Parametric tests were used only after testing data distribution for normality.

## Results

### Inhalation-linked patterning of mitral/tufted cell membrane potential

We obtained whole-cell current clamp recordings from OB neurons while controlling inhalation artificially in anesthetized mice. Artificial inhalation allowed for precise and reproducible control of olfactory sampling and, critically, allowed for the precise alignment of neuron responses across trials and across cells (Fig. 1A). A separate tracheal tube allowed inhalation of ambient air or odorant to be controlled independent of ongoing respiration (Sobel and Tank, 1993). The principal experimental paradigm involved recording responses to inhalation of ambient air and then odorant at each of three inhalation frequencies: 1, 3 and 5 Hz. We recorded from 39 cells, including 90 odorant presentations, in which all three frequencies were tested. Fifteen of these could be classified as mitral or tufted cells after filling with sulforhodamine, based on soma size and the presence of a clear apical dendrite extending to the glomerular layer. One filled cell (out of 16) was not clearly a mitral or tufted cell; this cell was classified as a granule cell and had a higher input resistance (500 MΩ) and recording depth (475 μm) than all but two of the 38 remaining cells; these three cells were excluded from further analysis Based on this result we presume that the large majority of unidentified recordings were from either mitral or tufted cells.

Respiratory patterning is a characteristic feature of MT cell activity; we thus first characterized modulation of membrane potential and spontaneous spiking by inhalation of ambient air at 1 Hz, which allowed for averaging of multiple inhalation-triggered responses with minimal interference across successive inhalations (Cang and Isaacson, 2003). 1 Hz inhalation is near the minimum respiration frequency in awake mice (Wesson et al., 2008). We constructed ‘inhalation-triggered averages’ (ITAs) of membrane potential, aligned to inhalation onset, over multiple (15–115; median, 40) inhalations of ambient air per cell (Fig. 1B). Approximately half (45%; 17/36) of all cells showed significant inhalation-linked modulation of membrane potential as measured from ITAs (see Methods for analysis details) during 1 Hz inhalation. Most (15/17) inhalation-driven responses consisted of brief depolarizations, with latencies to peak depolarization ranging from 43 - 387 msec after peak inhalation pressure (mean ± s.d., 196 ± 92 msec) (Fig. 1C).

Earlier reports in anesthetized rodents have reported distinct respiratory patterning of MT cell subclasses defined by soma depth (Onoda and Mori, 1980; Griff et al., 2008; Fukunaga et al., 2012; Igarashi et al., 2012). To examine this in our artificial inhalation paradigm, we classified filled MT cells as superficial, middle, or deep, based on the location of their soma relative to the mitral cell layer (Fukunaga et al., 2012) (Fig. 1D); while these designations might correspond to the classical MT subtypes of superficial, middle, and internal tufted, and mitral cells, respectively, we did not attempt to apply this designation given the lack of additional morphological or physiological criteria. Finally, we likely did not record from the most superficial MT cells whose somata reside in the deep glomerular layer. In total, we identified 7 deep (22 cell-odor pairs), 4 middle (8 cell-odor pairs), and 4 superficial MT cells (12 cell-odor pairs), with the remainder (23 cells, 47 cell-odor pairs) unidentified.

Significant modulation of membrane potential by inhalation was present in 3 of 7 deep MT cells, 2 of 4 middle MT cells and 2 of 4 superficial MT cells. These cell depth classes showed overlapping temporal responses in terms of latency or phase relative to inhalation (Fig. 1C, E). We further investigated this relationship by including unidentified recordings in the dataset; because soma depth and axial recording depth were well correlated for successfully filled cells (Fig. 1F; r = -0. 82, n = 15), we used recording depth to nominally distinguish deep from superficial MT cells (deep, >350 µm; superficial, <275 µm). This analysis yielded 4 superficial and 8 deep MT recordings (plus 5 cells at intermediate depths) with significant modulation by inhalation (Fig. 1G). However, we again saw no difference in the time to peak depolarization across these two classes (superficial median, 179 msec; deep median, 185 msec; p = 0.80, Mann-Whitney test). We also measured inhalation-linked temporal pattering as a function of the phase of the respiratory period, to account for the entire pattern of modulation rather than simply latency to peak (Goldberg and Brown, 1969; Fukunaga et al., 2012; Youngstrom and Strowbridge, 2015). Using this approach, of filled cells, 5 of 7 deep MT cells and 4 of 8 middle or superficial MT cells were significantly modulated (p<0.05, Rayleigh test). Among all cells sorted by recording depth, significant modulation occurred in 5 superficial and 11 deep MT cells. However, in agreement with latency analysis, the range of preferred phases in the inhalation cycle was highly overlapping for superficial and deep cells (mean θ = 2.73 and 2.69 rad, respectively) (Fig. 1G, inset). Thus, in contrast to earlier reports in ketamine-anesthetized, freely-breathing mice (Fukunaga et al., 2012; Igarashi et al., 2012), we could not reliably distinguish superficial from deep MT cells based on the timing of their inhalation-evoked activity.

### Diversity of temporal patterns elicited by odorant inhalation

Next we assessed odorant-evoked responses during artificial inhalation at 1 Hz using single odorants known to evoke input to the dorsal OB, presented at low-to moderate concentrations (0.1% - 2% s.v.) (Fig. 2). We restricted our analysis only to odorants effective at eliciting a change in spiking or in Vm when presented (see Methods). Nearly all cell-odor pairs showed significant inhalation-linked patterning of membrane potential by effective odorants (35 of 36 cells and 82 of 90 cell-odor pairs). In contrast to air inhalation, however, odorants evoked a diversity of response types, consistent with previous reports (Fig. 2A-C). Odorant-evoked ITAs ranged from a simple depolarization or hyperpolarization to more complex multiphasic responses with hyperpolarizing and depolarizing components (Fig. 2A-C). ITA shape – including the relative magnitude of hyperpolarization or depolarization and the timing of peak depolarization – could vary within the same cell for different odorants (Fig. 2A) and for different concentrations of the same odorant (Fig. 2B). While ITA latency sometimes decreased with increasing odorant concentration, concentration did not consistently impact latency or amplitude: of six MT cells (8 cell-odor pairs) tested at multiple concentrations spanning at least a 2- to 10-fold range (Fig. 2D), peak ITA Vm amplitude and latency at the low and high concentrations were not significantly different (median amplitude at 1 Hz = 3.90 and 2.89 mV for low and high concentrations; p=0.44, Wilcoxon signed rank test, n = 8 cell-odor pairs; median latency to peak depolarization at 1 Hz = 286 and 219 msec for low and high concentrations; p=0.31, Wilcoxon signed rank test, n = 6 cell-odor pairs).

We sorted odorant-evoked ITA patterns across all cell-odor pairs according to their predominant response polarity (i.e., depolarizing or hyperpolarizing) and, secondarily, by latency to peak deviation from rest (Fig. 2E). For predominately depolarizing (excitatory) responses (n=59 cell-odor pairs), latencies to peak depolarization ranged from 58 - 378 msec (median ± s.d., 191 ± 76 msec); predominately hyperpolarizing responses (n=23 cell-odor pairs) had latencies to trough ranging from 61 – 477 msec (median ± s.d., 127 ± 99 msec). Inhalation-triggered changes in spike rate – including spike suppression – showed dynamics closely matching those of the Vm ITAs (Fig. 2F).

Recent studies have provided evidence that superficial versus deep MT cells show distinct respiratory patterning and differences in the degree to which they are shaped by inhibition during odorant inhalation (Igarashi et al., 2012; Adam et al., 2014; Economo et al., 2016). However, in our recordings, neither the prominence of hyperpolarizing responses nor the latency to peak excitation varied systematically as a function of morphologically-identified cell type (Fig. 2E) or as a function of recording depth (Fig. 2G). For example, identified superficial MT cells could show clear hyperpolarizing components in their ITA (e.g., Fig. 2A), and had similar latencies to peak depolarization as identified mitral cells (superficial MTs: median = 190.7 msec, n = 9 cell-odor pairs; deep MTs: 224.3 msec, n = 17 cell-odor pairs; Mann-Whitney test, p = 0.056). This analysis suggests that the inhalation-linked dynamics of odorant-evoked responses are diverse and have the potential to be shaped by inhibition in both superficial and deep mitral and tufted cells (Yamada et al., 2017).

### Effect of inhalation frequency on inhalation-linked response patterns

We next investigated how MT responses change with repeated sampling of odorant at different frequencies. Sustained, high-frequency sniffing of odorant is a hallmark of active odor investigation and of exploratory behavior in general (Welker, 1964; Wesson et al., 2008), although the impact of this sampling on odor representations across MT cells remains unclear. We analyzed the impact of sniff frequency on inhalation-driven patterning, spike timing, and mean membrane potential.

Inhalation-linked patterning of membrane potential or MT spiking persisted at higher inhalation frequencies, although in a lower proportion of cells, with 43 of 90 cell-odor pairs showing significant inhalation patterning at 5 Hz compared with 82 of 90 cell-odor pairs at 1 Hz. Increasing inhalation frequency altered and simplified the temporal pattern of the inhalation-linked response for individual cell-odor pairs by shortening the period over which responses could evolve before being impacted by the next inhalation. Figure 3A shows an example recording from an identified deep MT cell during inhalation of the same odorant at 1, 3 and 5 Hz. This cell responded with a simple depolarization and spike burst linked to inhalation whose latency to peak remains relatively consistent at each frequency but whose duration shortens. Likewise, in a second example, the deep MT cell shown in Figure 3B had an ITA response consisting of a multi-phasic hyperpolarization-depolarization sequence at 1 Hz, which morphed into a simple sinusoidal fluctuation in membrane potential - and a single coherent spike burst - for 3 and 5 Hz inhalations.

**Figure 3.**
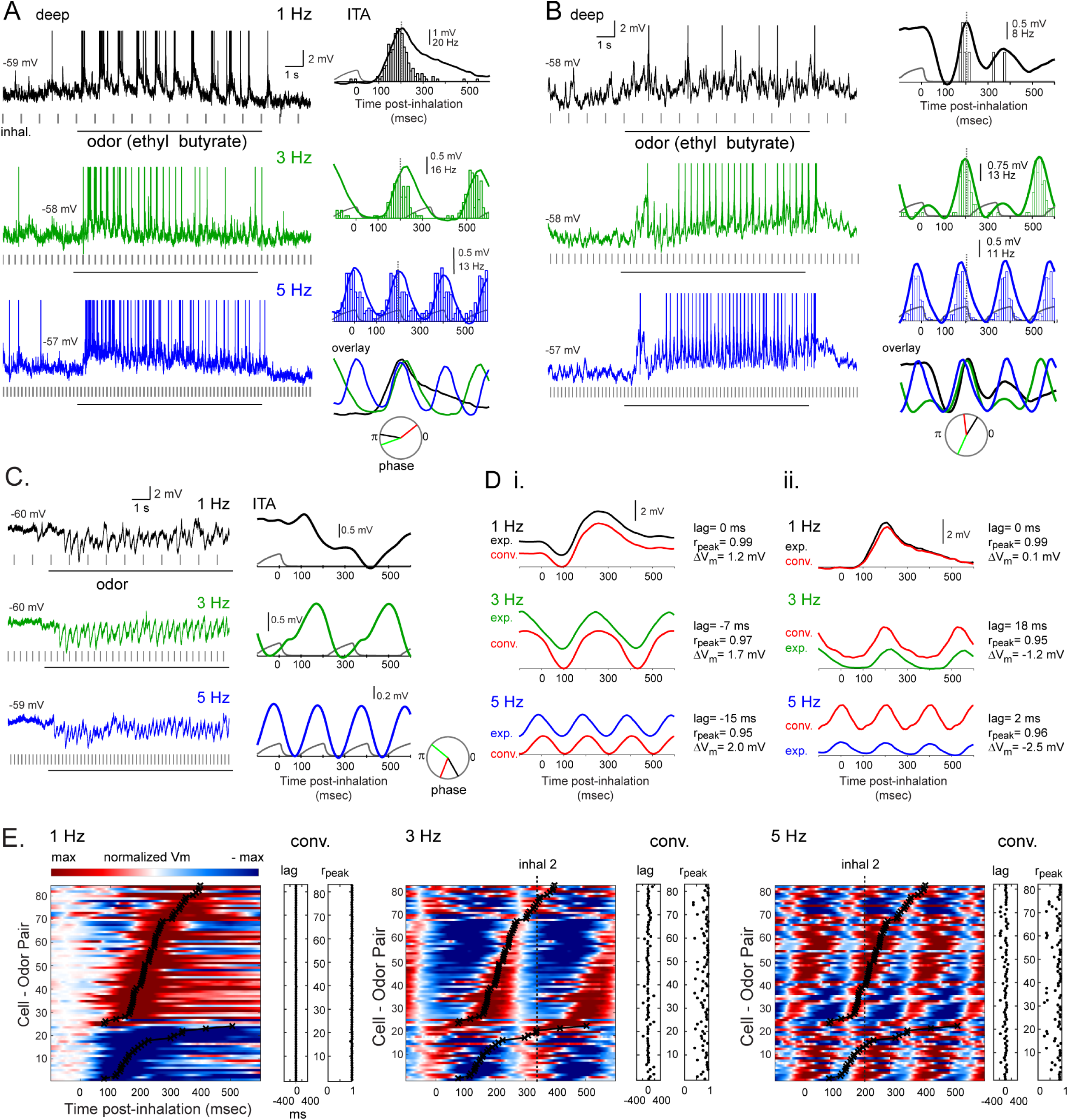
Inhalation-linked temporal patterning of odorant responses persists across frequency. A. Raw V_m_ traces (left) and odorant-evoked ITA of V_m_ and spiking (right) for a deep MT cell during 1, 3 and 5 Hz inhalations. Depolarization and spike burst duration shortens at higher sniff frequencies, but peak time remains consistent. Dashed line indicates time of peak depolarization in the 1 Hz ITA, for comparison across frequencies. ITAs for each frequency are overlaid at lower right for visual comparison. Circular plot shows phase of ITA for each frequency. B. Second example showing odorant-evoked ITA of V_m_ and spiking for a deep MT cell during 1, 3 and 5 Hz inhalations, with a biphasic hyperpolarization/depolarization sequence morphing to a simple sinusoidal modulation at 3 and 5 Hz, along with an emergence of spiking. C. Raw Vm traces (left) and ITA (right) from a deep MT cell showing the emergence of sinusoidal inhalation-linked patterning of Vm from a purely hyperpolarizing response to odorant inhalation at 1 Hz. D. Comparison of recorded (‘exp’) and predicted (red, ‘conv.’) Vm ITAs for the cells shown in panels A (i) and B (ii). Predicted ITAs were generated from convolution of the recorded 1 Hz ITAs with the inhalation event times at 1, 3 and 5 Hz. See Text for details and for definition of ‘lag’, ‘rpeak’ and ‘ΔVm’. In both cells, ITA shape is similar for recorded and predicted traces, but convolution does not predict mean Vm well, with ΔVm increasing in magnitude at higher frequencies and varying in polarity for both cells. E. Pseudocolor plots of odorant-evoked Vm ITAs for all responsive cell-odor pairs at 1, 3 and 5 Hz, sorted by latency to peak depolarization or hyperpolarization at 1 Hz. 1 Hz plot is identical to that from Fig. 2D. Black plot overlay indicates peak time for 1 Hz inhalation, for comparison across frequencies. ITAs are low-pass filtered at 50 Hz, then scaled as in previous figures (max to -max deviation from baseline, with baseline set to the mean pre-inhalation Vm. ITAs at each frequency were scaled independently. Dashed vertical line (‘inhal. 2’) indicates peak time of next inhalation. At 5 Hz, in approximately 40% of cells depolarization begins after the next inhalation has occurred. The lag and rpeak values comparing the recorded and convolved (predicted) ITA for each cell-odor pair are shown to the right of each pseudocolor plot, with each point corresponding to the same cell-odor pair to its left. See Text for details.

Another notable feature of MT responses at higher frequency was that the temporal dynamics of inhalation-linked membrane potential changes and spike bursts could appear identical for cell-odor pairs that showed opposite-polarity responses at 1 Hz inhalation. For example, at 1 Hz, the mitral cell in Figure 3B responds with an initial hyperpolarization which is absent in the mitral cell shown in Figure 3A, while a third mitral cell (Fig. 3C) shows a purely hyperpolarizing response to inhalation. Nonetheless, the 5 Hz ITA for all three cells is sinusoidal with similar latencies to peak depolarization. These observations suggest that similar inhalation-linked patterning of MT cell responses can arise from excitatory as well as inhibitory circuits, and that the relative strength of excitatory and inhibitory pathways in MT responses at low frequency does not simply map to inhalation-linked patterning at higher frequency.

Inhalation-linked responses did not uniformly compress in time as inhalation frequency increased, as might be expected if MT response timing represented odor information with respect to respiratory phase (Chaput, 1986; Smear et al., 2011). The preferred phase of individual MT cell responses shifted with inhalation frequency (Figs. 3A-C), with phase shifts varying substantially across cells (mean phase shift from 1 to 3 Hz = 0.012 ± 0.35 rad; from 1 to 5 Hz = -0.0012 ± 0.38 rad). Instead, changes were more consistent with an absolute latency-based response pattern which was repeated with each inhalation and summed with the ongoing responses preceding it. To visualize this we compared ITAs of Vm recorded at 3 and 5 Hz with synthetic ITAs constructed by convolving the 1 Hz ITA with inhalation timing at these two frequencies (see Methods). Example of this comparison are shown for two MT cells in Fig. 3D. In these two examples, linear convolution accurately predicts the shortening of inhalation-triggered depolarizations, and predicts the timing of peak depolarization with only modest differences from the timing of the recorded responses.

Linear convolution of 1 Hz ITAs predicted peak depolarization times with relatively high accuracy across cell-odor pairs. To quantify this we measured the lag of the peak cross-correlation between experimental and convolved ITAs (see Methods); across the population, the mean lag remained near zero at 1, 3 and 5 Hz, although variance across cell-odor pairs increased at 3 and 5 Hz (mean ± s.d. for 1, 3 and 5 Hz = -1 ± 3 msec, 7 ± 67 msec, and 1 ± 66 msec, respectively; Figure 3D, E). Increasing inhalation frequency had a larger - but still relatively modest - nonlinear effect on the overall temporal pattern of the response, as measured by the peak correlation coefficient obtained in the cross-correlation analysis (Fig. 3E). Peak cross-correlations (rpeak) decreased from a mean of 0.99 (± 0.01) at 1 Hz to 0.81 (± 0.17) and 0.80 (± 0.21) at 3 and 5 Hz, respectively. Notably, for MT cell-odor pairs with slower excitatory responses (i.e., longer latencies to peak depolarization), high-frequency sniffing led to responses whose times of peak depolarization lagged a full inhalation cycle. This was the case for approximately 50% of cell-odor pairs tested at 5 Hz (Fig. 3E). In these cases, during the 5 Hz inhalation bout the latency of the response relative to inhalation onset was ambiguous without prior knowledge of the particular inhalation by which it was driven.

### Inhalation frequency has diverse and nonlinear effects on MT cell excitability

While linear convolution of MT cell responses at 1 Hz was a relatively good predictor of inhalation-linked temporal patterning of membrane potential at higher frequency, it did not accurately predict changes in mean membrane potential averaged over one or more inhalation cycles. For example, for the two MT cells shown in Fig. 3D, the mean Vm (over 1 second) of the experimentally-recorded ITA differs from the convolved prediction by several mV (‘ΔVm’, Fig. 3D). This ΔVm value varied in polarity for different cell-odor pairs (compare left and right columns in Fig. 3D), and, across all cell-odor pairs, increased in variance at higher frequencies (mean ± s.d. for 1, 3 and 5 Hz = 1.23 ± 1.87, 1.49 ± 3.38, and 1.51 ± 5.00 mV). Non-zero ΔVm values could not be explained by drift in pipette offset or other experimental errors in baseline Vm measurements, as recalculating ΔVm after correcting for occasional shifts in spike threshold Vm at 3 and 5 Hz relative to those at 1 Hz gave near-identical results (ΔVm for 3 Hz = 0.93 ± 6.2; 5 Hz = 0.97 ± 7.1 mV, with the standard deviation of the spike threshold measurement at 1 Hz = 2.37mV).This analysis suggests that inhalation frequency has nonlinear effects on overall MT cell excitability that do not arise solely from summation of unitary inhalation-driven responses.

Consistent with this analysis, we found that inhalation frequency had heterogeneous and nonlinear effects on overall MT cell excitability as reflected in mean or peak spike rate across an inhalation cycle. Across all recorded cell-odor pairs, normalized peak inhalation-evoked MT firing rates remained high over the course of the 10-second odorant presentation and were similar for 1, 3 and 5 Hz inhalations (Figure 4A). However, for *individual* cell-odor pairs the effect of frequency on spike rate was diverse, with sustained sampling at higher frequencies leading to both increases and decreases in mean rate per 1-sec window or peak spike rate per 200 msec, relative to those at 1 Hz (for 3 Hz, mean and peak rate change = -0.48 ± 3.5 Hz and -3.0 ± 20 Hz, respectively; for 5 Hz, mean and peak rate change = -0.32 ± 4.6 and -2.3 ± 26 Hz ) (Figure 4B).

**Figure 4.**
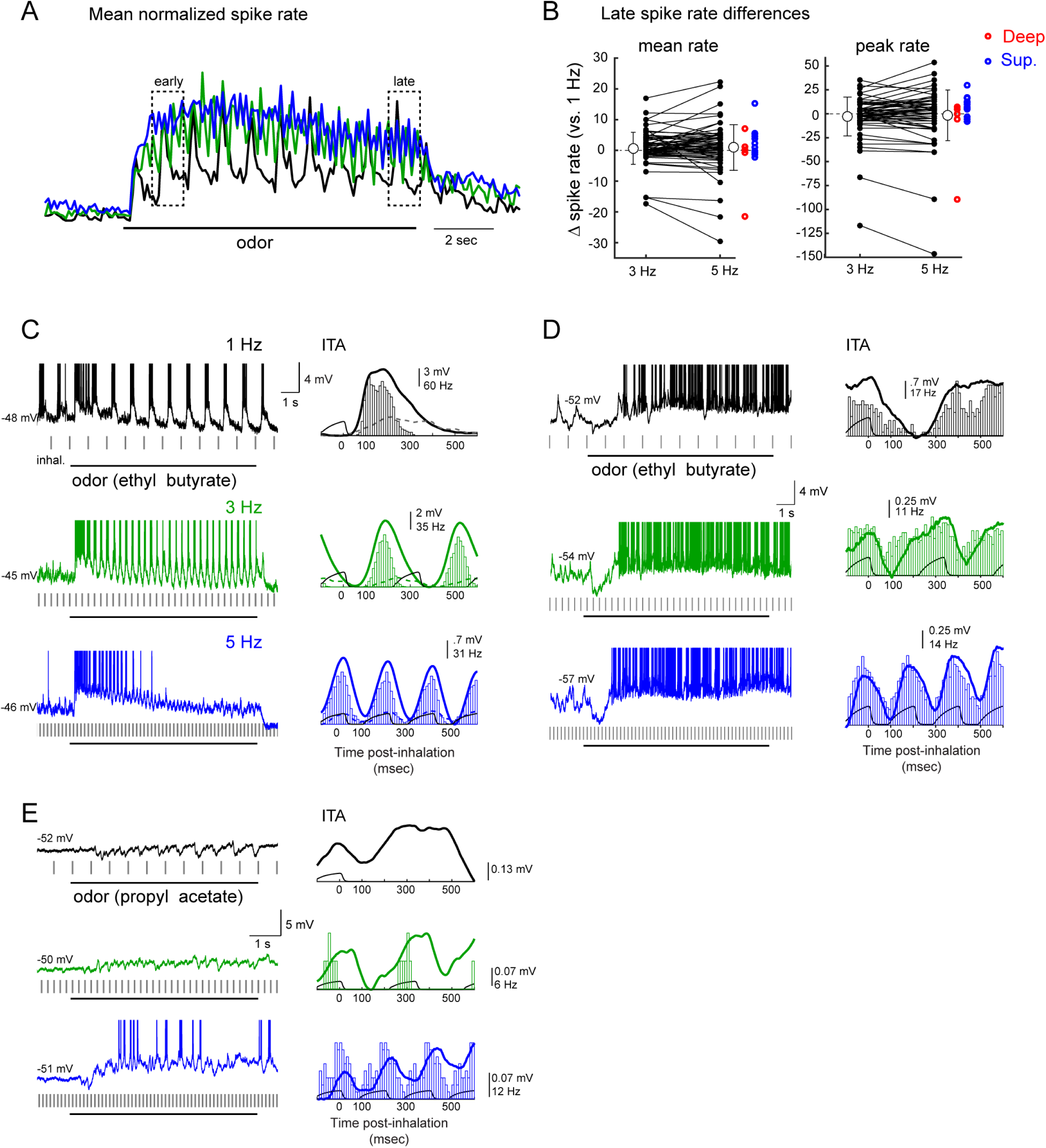
Sustained odorant sampling causes diverse changes in MT cell excitability that vary across frequency and cell identity. A. Traces showing normalized spike rate averaged across all responsive MT cell-odor pairs (n = 59) for 1, 3 and 5 Hz inhalations. The spike rate trace for each cell-odor pair (see Text for details) was normalized to its own maximum at a given frequency and then averaged with all other cell-odor pairs. Responses across the population show modest changes in rate over the 10-sec odor presentation which are similar across frequencies. Dashed boxes indicates time windows used to compute ‘early’ and ‘late’ spike rates in subsequent analyses. B. Diversity of effects of sustained sampling at 3 and 5 Hz across individual MT cell-odor pairs. Plots show difference in mean spike rate in the ‘late’ response (9 - 10 s after the start of odorant presentation), relative to that at 1 Hz for each cell-odor pair, for 3 Hz and 5 Hz inhalations. Both mean rate and peak rates over the 1-sec window are plotted. Note the high variance from 1 Hz rates, and that the mean difference from 1 Hz rates (open circles) is zero across all cell-odor pairs. Grey lines connect points from the same cell-odor pair. Error bars indicate s.e.m. Individual points for 5 Hz are duplicated to the right from each of the superficial (red) and deep (blue) MT cells. C-E. Examples from individual MT cells showing diversity of effects of inhalation frequency on overall excitability (spike rate). Sustained sampling can lead to adaptation of spiking (C) or an emergence of spikes after an initial hyperpolarization (D, E) in different cell-odor pairs.

For example, the MT cell in Fig. 4C shows a reduction in spike rate from early to late in the odor presentation at 1 Hz inhalation (see Fig. 4A for early and late time-windows) which transitions to complete adaptation at 5 Hz. In contrast, in the cell in Fig. 4D, spiking emerges after approximately 2 seconds of odorant inhalation at 1 Hz, with ongoing spiking interrupted by an inhalation-linked hyperpolarization; this spiking phase of the response emerges sooner and reaches higher rates at 3 and 5 Hz inhalations. In still other cells, increasing inhalation frequency led to subthreshold responses giving way to spiking responses (e.g., Fig. 3B), or resulted in responses reversing polarity from purely inhibitory at 1 Hz to excitatory at 5 Hz (Figure 4E). The effect of inhalation frequency on MT spiking was not predicted from the linear convolution of membrane potential ITAs at 1 Hz: we found a negligible correlation between the change in mean spike rate and the change in the mean Vm of the convolved ITA as inhalation frequency increased to 3 Hz (r for change in mean and peak ‘late’ spike rate from 1 Hz to 3 Hz = 0.06 and -0.08) and 5 Hz (r for mean and peak ‘late’ spike rate, 1 Hz to 5 Hz = 0.14 and 0.07, respectively). Latency of the ITA at 1 Hz also failed to predict the impact of inhalation frequency on MT spiking, with no significant correlation between peak ITA latency and change in mean or peak spike rate from 1 Hz to 5 Hz (r = 0.1 and 0.02 for mean and peak late rates, respectively). We also did not find any significant difference between the effect of inhalation frequency on mean or peak spike rate for superficial versus deep MT cells (p = 0.39 and p = 0.24 for mean and peak spike rate differences from 1 to 3 Hz and p = 0.10 and p = 0.13 for mean and peak spike rate differences from 1 to 5 Hz, Mann-Whitney test, superficial cells n = 8, deep cells n = 19) (Fig. 4B, red and blue points). We also tested for differences between ‘silent’ (< 1 Hz pre-odor firing rate) (Kollo et al., 2014) versus spontaneously-firing MT cells. High pre-odor firing rate cell-odor pairs (>1 Hz, n =13) had significantly different odor-evoked firing rates from low (<1 Hz, n=57) pre-odor firing rate cell-odor pairs (H(1,189)=17.0, p =0.00004; Scheirer-Ray-Hare extension of the Kruskal-Wallis test) (Sokal and Rohlf, 1995) and the effect of inhalation frequency on both groups was also significant (H(2,189)=20.9, p=0.00004), but the interaction between pre-odor firing rate and effect of frequency was not significant (H(2,189)=0.623, p=0.732). Thus, while pre-odor firing rate did impact mean odor-evoked firing rate, the effect of inhalation frequency was not different between high and low pre-odor firing rate cell odor pairs. Overall, these analyses indicate that inhalation frequency can profoundly shape MT excitability via nonlinear processes that vary across different MT cells.

### Sustained high-frequency sampling decorrelates odor representations

To further examine nonlinear effects of odorant sampling, we compared inhalation-evoked responses over the time-course of the 10-sec odorant presentation. To evaluate time-dependent changes in inhalation-linked patterning of Vm, we compared ITA waveforms averaged from early to late in the odorant presentation. The ITA showed relatively modest changes in shape at any of the frequencies (mean rpeak and lags for 82 cell-odor pairs, early versus late ITA = 0.77 ± 0.19 at -3.1 ± 144 msec, 0.72 ± 0.19 at -4.4 ± 146 msec and 0.66 ± 0.18 at 4.6 ± 182 msec at 1, 3 and 5 Hz, respectively). Thus, inhalation-linked temporal patterning of membrane potential remains relatively consistent across a prolonged sampling bout.

In contrast, as is clear from the examples in Figure 4, MT cell firing rates could change dramatically over time (e.g., Fig. 4C, D). While sustained sampling altered mean MT firing rates in some cells even for 1 Hz inhalations, this effect was more pronounced - and less uniform - at 3 and 5 Hz inhalation frequencies, with the linear correlation (r) between ‘early’ versus ‘late’ firing rates decreasing from 0.70 at 1 Hz to 0.46 and 0.42 at 3 and 5 Hz, respectively (n = 59 cell-odor pairs) (Fig. 5A). Similar results were obtained using peak inhalation-linked firing rate as a response measure (r values of 0.87, 0.48, and 0.38 for 1, 3 and 5 Hz, respectively). Thus, odorant-evoked spiking responses of MT cells change substantially across sustained sampling during high-frequency inhalations as compared to low-frequency inhalations. These changes in excitability typically developed over the first 1 – 2 seconds of odorant stimulation – or multiple cycles of inhalation, as we quantify below.

**Figure 5.**
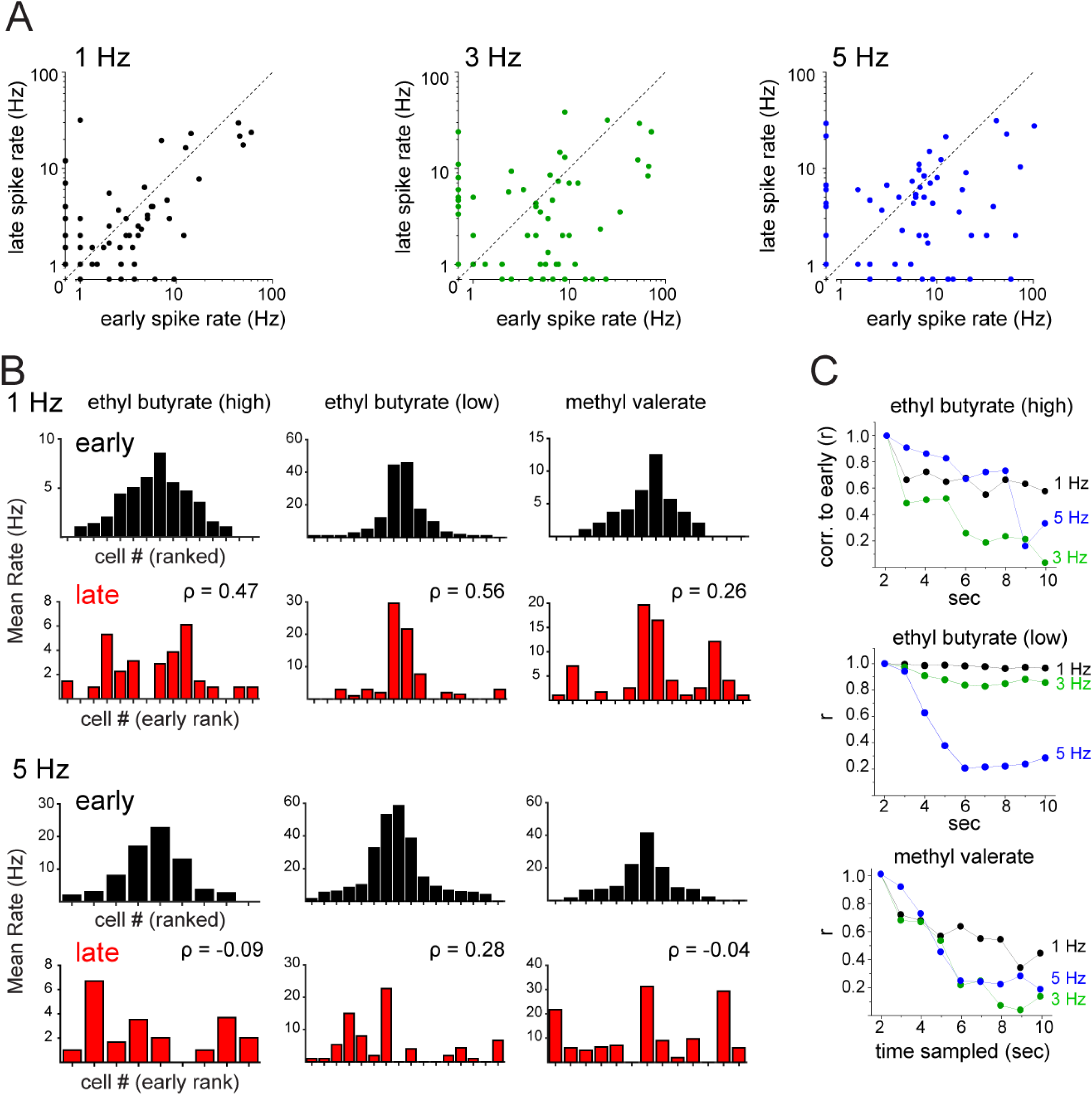
Sustained odorant sampling at higher inhalation frequencies decorrelates MT population response patterns. A. Mean spike rate during the early period of the odor presentation (seconds 1–2) plotted against mean spike rate during the late period (seconds 9–10) for responsive, spiking MT cell-odor pairs. B. Bar plots of mean firing rates during the early (black) and late (red) odor periods of trials with 1 Hz (top two rows) or 5 Hz (bottom two rows) inhalations for MT cells responding with spiking. Populations of responsive MTs are shown together for higher-concentration ethyl butyrate (left column), lower-concentration ethyl butyrate (middle column), and methyl valerate (right column; see Text for details). Bars are ranked from the center outwards by the mean firing rate within an odor during the early period and the order kept the same for the late period plots for comparison. The rank order of firing rates differ between the early and late periods, but these differ more at higher frequencies. The rank order correlation between early and late periods (Spearman’s ρ) is substantially lower at 5 Hz then at 1 Hz for all three odors. C. Linear correlation over time of mean firing rates of the same three virtual MT cell populations. Each point represents the correlation (Pearson’s r) of the mean firing rates between the early period (seconds 1–2) and each subsequent second. Correlations decrease at later time points for all frequencies, but decrease more, and more rapidly, at 3 and 5 Hz than at 1 Hz.

The diverse effects of repeated odorant sampling imply that odor representations involving patterns of activity across multiple MT cells would change during the course of a sampling bout. We simulated such an effect by compiling virtual populations of MT cells from multiple recordings of MT cells responding to the same odorant at a similar concentration, as has been done to characterize MT population coding in previous studies (Friedrich and Laurent, 2001; Barnes et al., 2008). We first compared MT population response patterns at the beginning and end of odorant presentation using rank correlation of mean spike rates (Fig. 5B). This analysis was performed for one odorant (ethyl butyrate) at two intensities and a second odorant (methyl valerate) at a single intensity. With all three virtual odor representations, the rank order of MT cell responses changed modestly during repeated 1 Hz inhalations but changed much more during 5 Hz inhalations. Likewise, using mean firing rate (per 1-sec window) as a response measure and linear correlation to compare population responses, virtual odor representations remained relatively stable throughout the odor presentation period at 1 Hz inhalations, but diverged from their initial pattern during 3 and 5 Hz inhalations (Fig. 5C). This analysis indicates that repeated sampling of odorants can dramatically reformat odor representations and that this reformatting is amplified at higher inhalation frequencies.

### Naturalistic odorant sampling also leads to diverse effects on MT cell odor responses

While artificial inhalation allowed us to isolate the impact of a single parameter of odorant sampling (i.e., frequency) on MT cell odor representations, active sniffing during behavior is accompanied by changes in other parameters such as inhalation duration or amplitude (Courtiol et al., 2011), which might alter the strictly frequency-dependent effects observed with artificial inhalation. To test for this possibility and to verify that high-frequency sniffing has diverse effects on odor representations using naturalistic sampling patterns, we performed additional MT cell recordings using a ‘sniff playback’ approach developed previously (Cheung et al., 2009; Carey and Wachowiak, 2011).

First, we measured respiration parameters in awake, head-fixed mice by recording intranasal pressure directly (Wesson et al., 2008; Wachowiak et al., 2013). Data were compiled from approximately 140 minutes of total recording time, including 28800 respiration cycles (inhalation and exhalation) across 5 mice (see Methods). As reported previously for freely-breathing rats (Courtiol et al., 2011), there was a significant change in inhalation duration across frequency (p < 2.2×10^−308^, Kruskal-Wallis test of inhalation durations binned by whole frequencies) (Fig. 6A), although there was substantial variability in duration at a given frequency, and the magnitude of this change was relatively modest at frequencies greater than 3 Hz (slope of best fit to binned duration medians across frequency for 4–10 Hz, -11.8 msec/Hz). Similarly, inhalation flow rate, estimated from the slope of inhalation onset for individual sniffs (see Methods), was significantly different across frequencies (p=8.4×10^-38^, Kruskal-Wallis test) but highly variable within a frequency bin (Fig. 6A). Thus, other parameters of odorant sampling do change with sniff frequency but are variable across individual sniffs.

**Figure 6.**
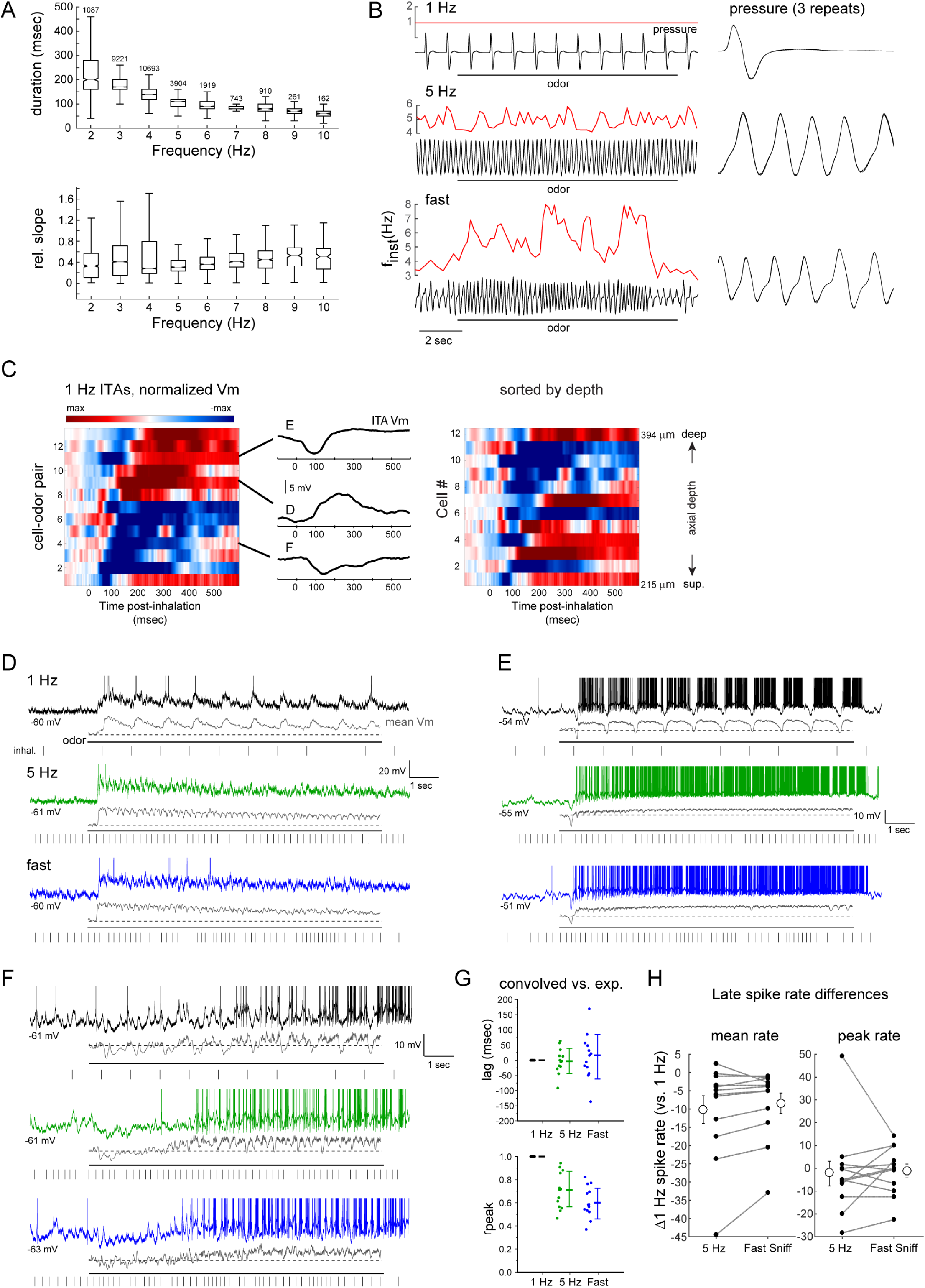
Heterogeneous effects of sniff frequency on MT cell excitability persist during playback of naturalistic active sniffing. A. Analysis of inhalation duration (top) and onset slope (bottom) as a function of instantaneous sniff frequency, measured from awake, head-fixed mice. See Text for details. Box and whisker plots show median, first and third quartiles of duration or onset slope and 1.5 times the interquartile range for each frequency bin. Number of sniffs measured is shown for each bin. B. Pressure waveforms used for the sniff playback recordings, with instantaneous frequency (finst) shown for the two active-sniff waveforms (‘5 Hz’) and (‘fast sniff’). See Text for details. Black trace shows pressure output from the sniff playback device; instantaneous frequency is shown in red above each trace. Right: Expanded view of a portion of the playback pressure, repeated three times for each playback trace and overlaid. Inhalation is up in all cases. C. Left: Pseudocolored time-course of odorant-evoked ITA for 12 cell-odor pairs using playback of the 1 Hz sniff, aligned to inhalation peak. Middle: ITA traces for the three example cells shown in (D, E and F), plotted relative to the time of peak inhalation. Right: Same ITAs for unique cells (n = 13), sorted by recording depth; numbers indicate vertical pipette depth from dorsal bulb surface. D – F. Example traces showing responses in three MT cells to playback of the 1 Hz, 5 Hz and fast sniff waveforms. For each frequency, top trace shows raw Vm (with spikes clipped), with the mean spike-filtered Vm averaged from 3 - 5 repeated trials shown below. Rasters below each trace indicate inhalation peak times. G. Comparison of lag (top) and rpeak (bottom) values comparing the recorded and convolved (predicted) ITAs for all cells (see Text for details) for 1 Hz, 5 Hz and fast sniff playback traces. Each point is one cell; mean and s.d. are shown to the right of data points. H. Diverse effects of sustained odorant sampling during active sniffing frequencies, compared to 1 Hz sniffing across individual MT cell-odor pairs. Plots show difference in mean (left) or peak (right) spike rate in the ‘late’ period of the response (8 - 9 s after start of odorant presentation), relative to that at 1 Hz for each cell-odor pair, for the 5 Hz and ‘fast sniff’ playback traces. Note the high variance from 1 Hz rates. Grey lines connect points from the same cell-odor pair. Error bars indicate s.d.

To test the impact of naturalistic odorant sampling at different frequencies, we generated prototypical command traces from epochs of intranasal pressure measurements which spanned and extended the range of the pulsed artificial inhalation experiments and used these to control odorant sampling (see Methods; traces are shown in Fig. 6B). We recorded odorant-evoked responses from 12 presumptive MT cells using each of these three traces (see Methods). As with pulsed inhalation, individual ‘sniffs’ of odorant repeated at 1 Hz evoked diverse inhalation-triggered responses with latencies to peak depolarization ranging from 244 – 421 msec after the peak of inhalation (Fig. 6C). Longer-latency depolarizations were typically preceded by a hyperpolarization. There was no obvious relationship between recording depth and the presence of hyperpolarizing components or latency to peak depolarization (Figure 6C).

Figures 6D-F show example traces of responses from three MT cells to odorant sampling with the 1 Hz, 5 Hz and ‘fast sniff’ sequences. Inhalation-linked modulation of membrane potential persisted even during the high-frequency sniff, as was apparent from multi-trial averages of spike-filtered Vm traces. In addition, as with pulsed artificial inhalation, the timing of inhalation-linked Vm changes during sniff playback was generally well-predicted by convolution of 1 Hz sniff responses, albeit with somewhat higher variability. For example, the lag of the inhalation-triggered average Vm during 5 Hz sniff playback relative to the convolved ITA waveform was short relative to the length of a sniff cycle and centered around zero (mean ± s.d. r_peak_ for 1 Hz = 0.99 ± 8×10^-5^, 5 Hz = 0.70 ± 0.16; fast sniff = 0.61 ± 0.13; Fig. 6G); the shape of the ITA waveform itself was less well predicted at 5 Hz sniff playback, with lower r_peak_ values than for pulsed inhalation. Assessing inhalation-linked timing for the high-frequency sniff playback was more difficult due to the variable inter-sniff intervals of the playback trace, but predicted response traces generated by convolution of 1 Hz ITAs with the sniff playback inhalation times nonetheless showed strong temporal correlations with the V_m_ recordings at short lag times (lag for 1 Hz = 0 ± 0, 5 Hz = -2.3 ± 42; fast sniff = 11.6 ± 74 msec, Fig. 6G).Sampling odorants with the two higher-frequency sniffing traces led to diverse effects on overall MT cell excitability, as reflected in changes in spike rate as well as mean V_m_. For example, the cell in Figure 6D responds to 1 Hz odorant inhalation with a slow depolarization and a spike burst that adapts after several inhalations, with a short-latency hyperpolarization and a more pronounced spike adaptation emerging at the higher sniff frequencies. In contrast, the two cells in Figures 6E and F, each of which have a short-latency hyperpolarizing ITA that suppresses spiking, show an increase in spike rate with successive sampling as well as an increase in spike rate at higher sniff frequencies.

Overall, the difference in mean Vm (ΔVm) between the convolved and predicted ITAs increased in variance as sniff playback frequencies increased (mean ± s.d. ΔVm for 1 Hz = 4.6 ± 5.3, 5 Hz = 5.5 ± 8.6; fast sniff = 5.6 ± 8.6 mV). Likewise, similar to the observations with pulsed inhalation, the effect of sustained high-frequency sniffing of odorant on mean or peak spike rate, relative to that for 1 Hz sniffs, varied substantially across cells. The mean firing rate during the ‘late’ period of odor sampling decreased relative to that for 1 Hz sniffs for both 5 Hz and ‘fast’ sniffing, (median difference = -5.43 Hz and -4.83 Hz, respectively, p = .002 and p = 0.009, respectively, n = 12 cell-odor pairs, Wilcoxon signed rank test), while peak firing rates during this time period did not show a significant difference across all cell-odor pairs (median difference for 5 Hz and ‘fast’ sniffing = -5.0 Hz and -2.92 Hz, p = 0.17 and p = 0.61, respectively, n = 12 cell-odor pairs, Wilcoxon signed rank test). More importantly, however, for both measures there was a high degree of variability in the change in spike rate across different cells (Fig. 6H). Also, as with pulsed inhalation, the change in firing rate for higher-frequency sniffing did not correlate with the predicted change in Vm obtained from convolution of the Vm ITA at 1 Hz: the correlation (Pearson’s r) between mean and peak firing rate change between 1 and 5 Hz during the ‘late’ period and the change in mean Vm predicted by convolution was -0.082 and 0.025, respectively, and was 0.015 and 0.11, respectively, for the same measure correlating firing rate change and predicted Vm change between 1 Hz and ‘fast’ sniffing (n = 12 cell-odor pairs for both). Overall these results suggest that diverse impacts of odor sampling frequency on different MT cells also occur during the high-frequency bouts of sniffing exhibited in awake mice.

### Bulk optogenetic activation of OSN inputs leads to uniform effects of input frequency

The diverse frequency-dependence of MT cell odorant-evoked spiking responses could arise from differences in MT cell intrinsic properties, differences in adaptation across OSN populations converging onto different glomeruli, or from odorant-specific patterns of synaptic excitation and inhibition that change over the course of sustained odorant sampling at different frequencies. To attempt to distinguish these mechanisms, we next activated OSN inputs optogenetically rather than by inhalation of odorant, using mice expressing ChR2 in all OSNs (Smear et al., 2011) and directing light onto the dorsal OB surface (see Methods). For each MT recording, light pulses (75 - 100 msec duration) were repeated at 1, 3 and 5 Hz using the same 10-sec protocol as for odorant stimulation and the same pulse duration across frequencies, but with no artificial inhalation (e.g., Fig. 7A, lower). This approach eliminated the contribution of peripheral adaptation of OSN inputs and uniformly evoked input to all dorsal OB glomeruli in the absence of any inhalation-or odorant-driven sensory input.

**Figure 7.**
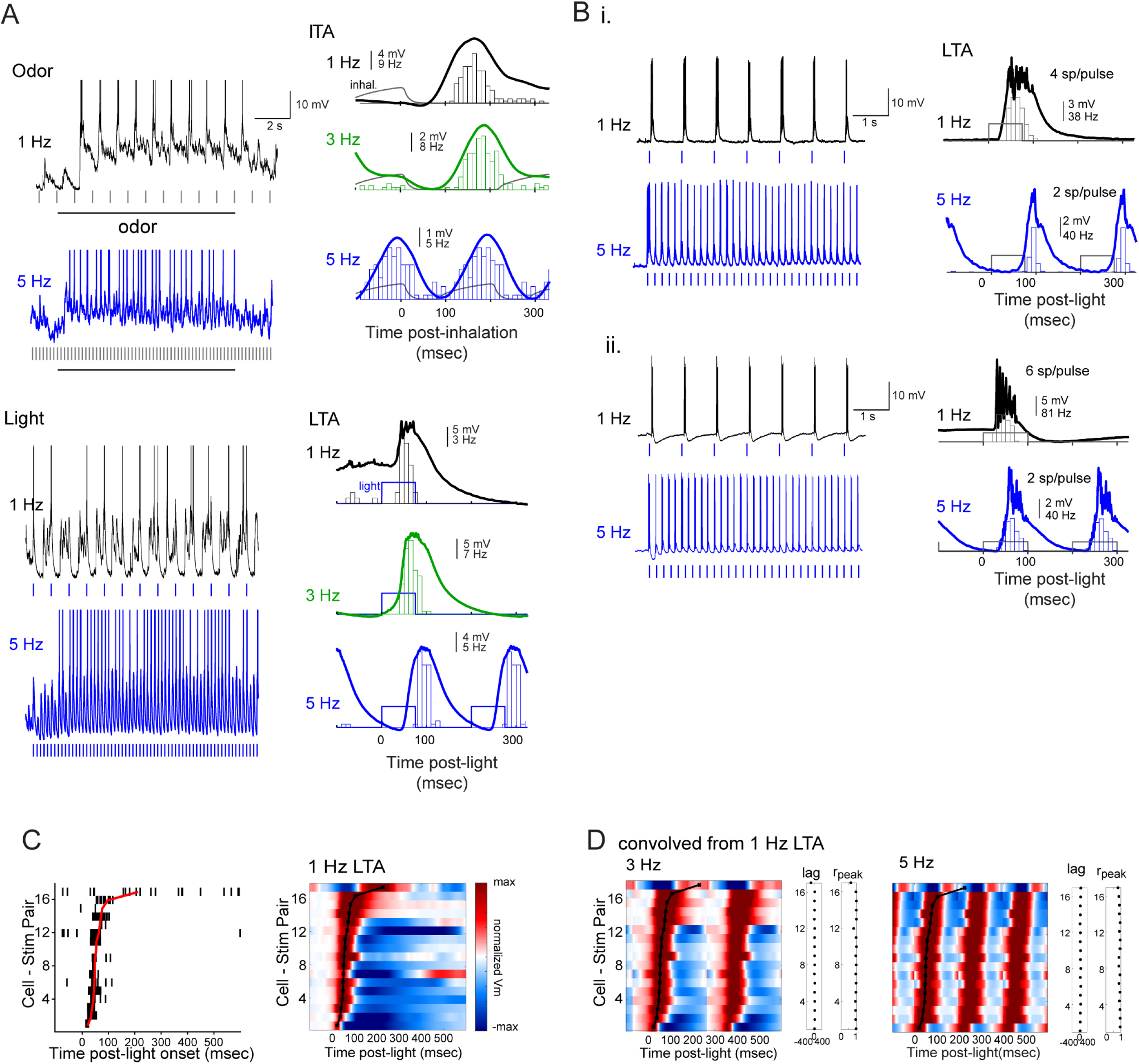
Bulk optogenetic activation of sensory inputs evokes robust, phasic MT cell responses across frequency. A. Upper: Raw V_m_ traces (left) and ITAs of V_m_ and spiking (right) for an unidentified MT cell in an OMP-ChR2 mouse during 1, 3 and 5 Hz inhalations. Lower: Response of same cell to light pulses at the same frequency, with raw V_m_ trace and light-triggered average (LTA; see Text for details). Blue marks indicate times of light pulses. Note shorter latency and duration of light-evoked compared to odorant-evoked responses. B. Examples from two additional MT cells showing raw Vm (left) and light-triggered average (LTA, right) for 1 and 5 Hz light pulses. While both MT cells reliably respond with spiking across frequency, the number of spikes per light pulse decreases from 1 to 5 Hz stimulation. C. Left: Light-triggered spike rasters for light-evoked responses for all MT cell-stimulus combinations, sorted by latency to peak depolarization (n = 11 MT cells; cells tested with multiple light intensities are plotted in separate rows). Red plot indicates time of peak depolarization, calculated from the light-triggered average (LTA) Vm trace. Right: Pseudocolored time-course of LTA of Vm for the same cell-stimulus pairs, sorted in the same order. Plots are normalized as in Fig. 2. Peak depolarization times are shown in black plot. Note that all but one MT cell (top row) showed an initial depolarization. Several MT cells only depolarized with no spiking in response to light. D. Pseudocolor plots of Vm LTAs predicted from convolution of the 1 Hz LTA at each frequency, presented in the same order as for the recorded 1 Hz LTAs shown in Fig. 6A. Black plots indicate recorded (experimental) 1 Hz LTA Vm peak times (identical to that from (C)). The lag and rpeak values comparing the recorded and predicted LTA for each cell-stimulus pair are shown to the right of each pseudocolor plot. See Text for details.

Optical stimulation of dorsal OB inputs evoked responses with a short-latency (mean ± s.d. = 70 ± 46 msec) depolarization in all recorded units (Fig. 7A-C) and, in most MT cells, a burst of 1 – 13 spikes per pulse. Light-evoked responses were typically shorter in latency and duration than responses to odorant inhalation in the same MT cell (Fig. 7A), although individual MT cells varied in the duration of their excitatory response and in the presence, magnitude and duration of subsequent hyperpolarizing response components (Fig. 7A-C). As with odorant stimulation, the temporal pattern of light-driven MT cell membrane potential changes at higher frequencies was well-predicted by linear convolution of the response at 1 Hz (Fig. 7D), with only a small increase in the latency to peak depolarization as frequency increased from 1 to 5 Hz for 11/17 cell-stim pairs (mean ± s.d. = 1 ± 7.5 msec, n = 17); overall response shape remained highly consistent across frequencies (mean r_peak_ at 3 and 5 Hz = 0.93 and 0.91, respectively).

The effects of increasing optical stimulation frequency on MT spiking were distinct and more uniform than with odorant inhalation. First, light-evoked spike rates averaged across the MT population decreased after the first light pulse at 3 and 5 Hz stimulation (Fig. 8A), whereas odorant-evoked firing rates increased gradually and then remained roughly constant (e.g., Fig. 4A). Second, there was less variation in the frequency-dependence of light-evoked excitation across individual MT cells, with 5 of 6 cells showing decreased light-evoked spiking at 3 Hz relative to 1 Hz stimulation and all MT cells showing a decrease at 5 Hz stimulation (Fig. 8B). Finally, after the first light pulse, the evolution of light-evoked spike rates over the course of the 10-second stimulus train was less pronounced and less variable across individual MT cells than for odorant-evoked responses: most MT cells showed little or no change in firing rate from the early to late stimulation epochs at 3 Hz, and a relatively linear decrease in firing rate at 5 Hz stimulation (r at 1, 3, and 5 Hz = 0.87, 0.54, and 0.77, respectively) (Fig. 8C). As a result, the rank order of light-evoked MT response magnitude (measured in spikes per flash) changed relatively little from the beginning to the end of the 10-sec stimulation, even for 5 Hz stimulation frequencies (Fig. 8D). This comparison suggests that the diverse and nonlinear effects of input frequency on odorant-evoked MT spiking responses are unique to odorant stimulation and do not arise solely from intrinsic differences among MT cells or differences in OSN adaptation. Instead, diverse frequency-dependence appears to emerge from odorant-specific network interactions within OB circuits, possibly driven by the combinatorial patterns of OSN input to OB glomeruli that are characteristic of odor representations.

**Figure 8.**
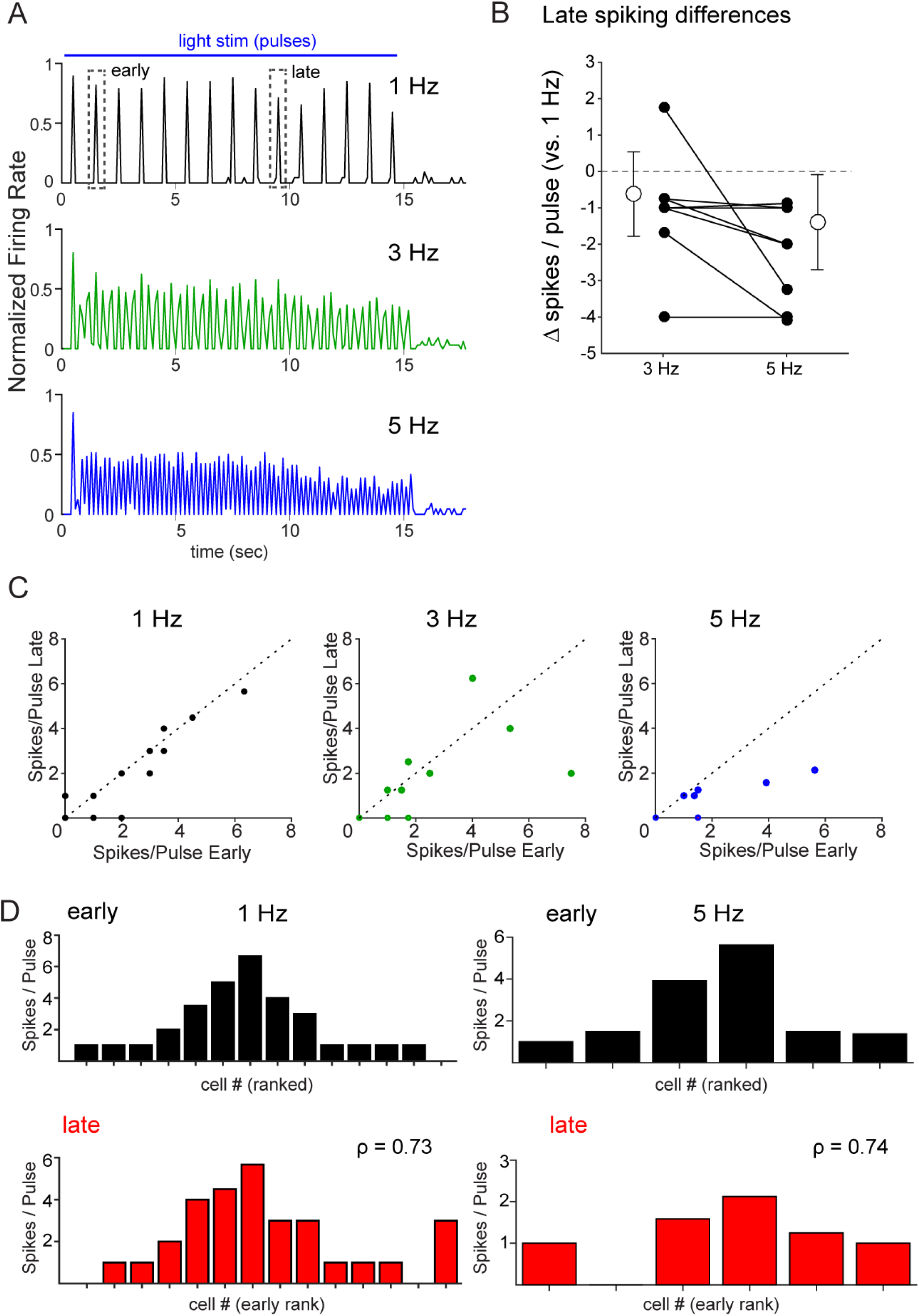
Uniform effects of input frequency in response to bulk optogenetic activation of sensory inputs. A. Traces showing normalized spike rate averaged across all responsive MT cell-stimulus pairs (n = 17) for 1, 3 and 5 Hz inhalations. The spike rate trace for each cell-stimulus pair was normalized to its own maximum at a given frequency and then averaged with all other pairs (see Text for details). Across the population, responses show modest changes in rate over the 15-sec light presentation that are similar across frequencies. Dashed boxes indicate time window used to compute ‘early’ and ‘late’ spiking in (B) and (C). B. Effects of sustained stimulation at 3 and 5 Hz across individual MT cell-stimulus pairs. Plots show difference in mean spikes per light pulse during seconds 9 - 10 of light stimulation relative to that at 1 Hz for each cell-stim pair, for 3 Hz and 5 Hz stimulation. Mean (open circles) and s.d. (error bars) of the 3 and 5 Hz groups are also plotted for each group. C. Mean spikes per pulse during the early period stimulation (seconds 1–2) is plotted against the mean spikes per pulse during the late period (seconds 9–10) for all MT cell-stimulus pairs (n = 17). The unity line is shown for comparison. Spikes per pulse remain consistent at 1 Hz and uniformly decrease at 5 Hz stimulation frequencies, with mixed effects across MT cells at 3 Hz. D. Histograms of mean spikes per pulse during the early (black) and late (red) stimulation periods of trials with 1 Hz (left) or 5 Hz (right) stimulation for MT cells responding with spiking, sorted by rank at 1 Hz, as in Fig. 5B. Spearman’s rank order correlation (ρ) of responsiveness between the early and late periods remains relatively high for both 1 Hz and 5 Hz stimulation, compared to that for odorant inhalation (see Fig. 5B).

## Discussion

Odor representations at the level of OB output are well known to change as a function of wakefulness, behavioral context, and experience (Bhalla and Bower, 1997; Kay and Laurent, 1999; Doucette and Restrepo, 2008; Kato et al., 2012; Yamada et al., 2017). However, disentangling effects of experience or neuromodulation from ‘bottom-up’ effects driven by changes in odor sampling is difficult, especially in awake animals which dynamically control odor sampling behavior. Here, by recording intracellular responses from MT cells while precisely controlling inhalation frequency, we were able to examine such bottom-up effects systematically and to test predictions from in vitro experiments on how inhalation frequency impacts MT cell responses. We found that inhalation frequency led to nonlinear changes in the overall excitability of MT cells which were not predicted from low-frequency responses, and that inhalation frequency had diverse effects across different cells. These results imply, first, that the effects of sustained and repeated odorant sampling are not uniform across all MT cells and, second, that repeated sampling can substantially reorganize MT cell odor representations - a process which is amplified at frequencies corresponding to active odor investigation.

Our results differ from those of several earlier studies examining the effects of odor sampling on MT responses recorded extracellularly. For example, Bathellier et al. compared MT spiking responses during 3 and 6 Hz artificial inhalation and found that the number of spikes per inhalation decreased by roughly half when inhalation frequency doubled, such that mean excitability stayed roughly constant (Bathellier et al., 2008). While we found this relationship was true for the MT cell population (e.g., Fig. 4A) there was large variation across individual cells, with some showing increases in spike rate and others showing decreases (e.g., Fig. 4B). Similarly, we found that MT spike rates – and mean membrane potential - at higher inhalation frequencies were poorly predicted from the inhalation-triggered response at 1 Hz, inconsistent with recent reports of a linear relationship between unitary MT spiking responses and odor stimulus profiles (Gupta et al., 2015). These changes occurred also during ‘playback’ of sniff bouts recorded from awake mice, strengthening the conclusion that inhalation frequency drives diverse and nonlinear changes in MT excitability in a bottom-up manner during active odor sampling.

Differences between these and previous results might be explained by a reliance on spiking for deriving unitary responses. For example, because of the inability of extracellular recordings to report inhibition directly, Gupta et al. derived ‘unitary’ MT response kernels from a large range of different temporal odorant profiles, whereas our whole-cell recordings allowed us to directly record subthreshold and inhibitory responses averaged over a single inhalation, which functionally defines the unitary response with respect to odorant sampling. The former approach may in effect encompass frequency-and time-dependent nonlinearities while deriving a unitary response kernel, leading to a similarly effective prediction when tested across different inhalation frequencies. It is also possible that this approach may effectively predict MT responses to rapidly-varying inhalation patterns, but not responses during the sustained bouts of inhalation at higher frequencies tested in the present study, in which nonlinear changes in spike rate emerged over seconds.

At the same time - and in agreement with the results of Gupta et al. (Gupta et al., 2015) - we found that, with respect to the temporal patterning of MT responses within an inhalation cycle, subthreshold response patterns at higher frequencies (3 and 5 Hz) were, in general, well predicted from those measured at 1 Hz. Simple linear convolution of 1 Hz response patterns could account for substantial transformations of the subthreshold response - including, for example, a shortening of response duration (e.g., Fig. 3A) and the emergence of regular sinusoidal responses from multiphasic or pure hyperpolarizing responses (e.g., Figs. 3B, 3F) - as inhalation frequency increased. Linear convolution analysis also predicted the ‘wrapping around’ of MT response latencies relative to inhalation for MT cells whose peak depolarization latencies exceeded that of an inhalation cycle period (e.g., Fig. 3D).

These results have implications for timing-based models of odor coding and processing. First, they indicate that the patterning of membrane potential by inhalations at different frequencies can be largely explained by linear summation of unitary (i.e., low-frequency) responses in absolute time, without requiring remapping of response timing to phase-based timing coordinates (Chaput, 1986; Khan et al., 2008; Shusterman et al., 2011). Indeed, the phase of inhalation-linked responses varied substantially - and differently for different cells - as frequency increased. The simplification of temporal response patterns at higher frequencies that arises from linear summation could impact coding strategies that rely on complex sequences of excitation and inhibition to encode odor information (Uchida et al., 2014). Second, they demonstrate that as inhalation frequency increases, there is increasing ambiguity in the latency of the MT response with respect to inhalation onset without prior knowledge of the particular inhalation by which it was driven. This ambiguity will impact the robustness of latency-based odor coding strategies (Brody and Hopfield, 2003; Spors et al., 2006; Shusterman et al., 2011) in the behaving animal, in which sniff frequencies can range from near 2 Hz up to 10 Hz or higher during active odor investigation (Kepecs et al., 2007; Wesson et al., 2008; Khan et al., 2012), because the identity of the earliest-responding MT cells will become less certain as inhalation frequency increases. Third, they suggest that coding strategies that rely on integrating activity across a particular time-window relative to inhalation - as has been proposed to occur in piriform cortical circuits (Poo and Isaacson, 2009; Franks et al., 2011) - should be robust across inhalation frequencies, as the timing of peak depolarization (and peak spike rate) was relatively invariant for a given MT cell-odor pair.

On a slower time-scale, our results suggest that increases in odor sampling frequency drive the emergence of nonlinear and tonic changes in MT cell excitability reflected in mean membrane potential and spike rate. Several cellular or circuit mechanisms might underlie odor-and cell-specific changes in tonic excitability. Reduced MT cell excitation at higher inhalation frequencies could result from adaptation of sensory neuron responses (Verhagen et al., 2007; Carey and Wachowiak, 2011), but also from an increase in the strength of feedforward or lateral inhibition onto MT cells via glomerular circuits (Gire and Schoppa, 2009; Shao et al., 2012; Shao et al., 2013; Liu et al., 2016). Circuit mechanisms underlying frequency-dependent increases in MT excitation might involve enhanced feedforward excitation by external tufted cells (Hayar et al., 2004; De Saint Jan et al., 2009) or disinhibition among glomerular or granule cell inhibitory circuits (Burton and Urban, 2015; Zak et al., 2015).

A crucial aspect of the frequency-dependent effect on MT excitability was its dependence on cell and odorant identity: whether higher frequencies led to increased or decreased excitation varied for different MT cells. Widespread activation of sensory inputs to dorsal OB glomeruli using optogenetic stimulation in the absence of inhalation was more uniform across MT cells and consisted primarily of a reduction in excitability. These more uniform frequency effects could be due to a greater degree of synchrony of OSN inputs to glomeruli or, more likely, to the homogeneous patterns of input to nearly all dorsal glomeruli. We hypothesize that, with odorant stimulation, changes in MT excitability depend on the frequency-dependence of the multiple OB circuits that shape the MT cell response, and that the relative contribution of these can vary as a function of differences in the combination of glomeruli receiving excitatory input and differences in the strength or temporal pattern of input across those glomeruli.

It is tempting to speculate that frequency-dependent enhancement or suppression of odorant-evoked spiking maps to different MT subtypes – for example superficial versus deep MT cells, whose evoked responses are thought to be differentially shaped by inhibitory OB circuits (Fukunaga et al., 2012; Igarashi et al., 2012; Phillips et al., 2012). However, MT soma depth did not predict the effect of increasing inhalation frequency on odorant-evoked excitation, nor did it correlate with the temporal pattern of the inhalation-linked response to either odorant or ambient air. Frequency effects were also not clearly correlated with spontaneous activity, response latency, peak amplitude or even polarity of the inhalation-linked membrane potential response at 1 Hz. Instead, the primary determinant of frequency-dependent changes in MT output may be the pattern of activity in glomeruli including and surrounding the parent glomerulus of the recorded cell. In this model, interglomerular circuits could differentially shape the strength of excitatory versus inhibitory inputs to a MT cell depending on the number and identity of nearby activated glomeruli (Economo et al., 2016). Future experiments that record MT cell responses across inhalation frequency simultaneous with imaging patterns of glomerular activation will be important to rigorously test this idea.

Regardless of the underlying circuit mechanisms, the diversity of inhalation frequency effects on MT excitability can have profound consequences for how sampling behavior can shape odor representations at the level of OB output. Our results strongly suggest that sustained high-frequency odor sampling leads to a reformatting of the MT population code for odor identity rather than a uniform change in gain or sharpening of response specificity. This reformatting may be analogous to that seen among zebrafish mitral cells during sustained odor presentation, in which mitral cell odor representations decorrelate to more reliably encode odor identity (Friedrich and Laurent, 2001). An important difference in the mouse, however, is that the magnitude of this reformatting depends on inhalation frequency and thus can be actively controlled by the animal’s sampling behavior.

The adaptive value of frequency-dependent reformatting of odor representations remains to be investigated. Rodents engage in sustained high-frequency sniffing when tracking odor trails (Khan et al., 2012), investigating novel stimuli (Macrides, 1975; Verhagen et al., 2007) and exploring (Welker, 1964), but not when performing learned odor discriminations (Wesson et al., 2009). In these contexts, high-frequency sniffing may decorrelate odor representations to facilitate fine odor discriminations, maximize sensitivity to changes in odor intensity, or support the efficient encoding of odor memories. Frequency-dependent reformatting might also facilitate the analysis of complex odor mixtures by altering the relative contribution of different mixture components to the population response, or by mediating a mixture-specific sequence of MT activity over the course of several seconds of high-frequency sniffing. While these functional hypotheses remain to be tested, the present results underscore the flexibility in odor representations at the level of the OB and highlight the degree to which sensory responses can be dynamically regulated during behavior through bottom-up mechanisms alone.

## Acknowledgments

The authors thank C. Zabawa and J. Ball for technical support, J. Fernandez, J. White and A. Schaefer for advice on recordings and Wachowiak lab members for helpful comments on the manuscript. D. Wesson collected the intranasal pressure data from awake mice. This work was supported by funding from NIH (DC06441 and DC013076).

